# The kinetochore proteins Ndc80 and Dsn1 are required for survival of postmitotic neurons

**DOI:** 10.64898/2026.06.29.735291

**Authors:** Guoli Zhao, Feng Tian, Qinglong Wang, Huyan Meng, Chen Ding, Richard T. Born, Zhigang He, Thomas Schwarz

## Abstract

Dsn1 and Ndc80 are essential proteins of the kinetochore complex and required for chromosome segregation in dividing cells and for regulating development and microtubule dynamics in postmitotic neurons. With conditional deletion of floxed alleles, we here show that Dsn1 and Ndc80 are also required for the viability of postmitotic neurons, both in cultures of hippocampal and cortical neurons and in vivo in the retina. Loss of these proteins triggers apoptosis, as indicated by caspase cleavage and an increase in nuclear DNA breakage. The pro-survival function of the kinetochore components is distinct from that which was previously demonstrated for regulation of neuronal synaptogenesis. The microtubule-binding domain of Ndc80 is required for the synaptogenic functions but Ndc80 lacking this domain can nonetheless rescue the viability of neurons from which Ndc80 has been deleted. Similarly, whereas the synaptogenic function involves regulation of microtubules in axons and dendrites, a nucleus-localized Dsn1 is sufficient to rescue the viability of neurons from which Dsn1 has been deleted. Thus, postmitotic neurons retain a nuclear requirement for components of the kinetochore in order to prevent apoptosis.

## Introduction

The kinetochore is a highly conserved, multi-protein complex recognized for its essential role in chromosome segregation during mitosis and meiosis. In these dividing cells it resides at centromeres and, by coupling the chromosomes to spindle microtubules, enables the sister chromosomes to separate and move to either pole of the cell [1–3]. Absent proper kinetochore function and regulation, cell divisions will undergo mitotic arrest, develop aneuploidy, and die. Emerging evidence, however, has begun to identify kinetochore functions beyond cell division. Recent studies across multiple species, including *C. elegans*, *Drosophila*, and mammals, have identified non-canonical, post-mitotic roles for kinetochore proteins in neuronal development. In *Drosophila*, kinetochore mutations have been shown to induce neurite overgrowth at the neuromuscular junction and in sensory dendrites [4, 5] and promote dendritic regeneration in sensory neurons [6]. Similarly, in *C. elegans*, these proteins are required to form proper neuronal circuits, including dendrite extension and axon fasciculation [7–9]. In mammalian systems, previous work from our lab demonstrated that kinetochore components regulate microtubule dynamics and the propensity of microtubules to invade nascent dendritic spines. Consequently, kinetochore components affect spine density *in vitro* and *in vivo*. These phenotypes identified a role for kinetochore proteins in fully developed, postpartum neurons that were long past their last cell division [4, 5].

The mitotic kinetochore is a large protein complex composed of distinct subcomplexes [1, 10]. The core of the complex, sometimes referred to as the KMN complex or outer kinetochore, consists of the three subcomplexes each named for a protein component: the Knl1, Mis12, and Ndc80 complexes. The Ndc80 complex, and Ndc80 protein in particular, interacts with the plus ends of microtubules, while the Mis12 complex connects the Ndc80 complex to CenpC and the inner kinetochore proteins that bind to centromeric DNA. The Knl complex is the linkage to many regulatory pathways [3, 11, 12]. Though the full nature of the neuronal kinetochore complex is unknown, studies in *C.elegans* and *Drosophila* have implicated each of these subcomplexes in postmitotic neuronal development of dendrites and axons [4–9].

Beyond these morphological abnormalities, we now demonstrate an essential role of kinetochore proteins for the survival of postmitotic neurons; kinetochore deficiency leads to progressive neuronal attrition by apoptosis. Moreover, the kinetochore functions required for neuronal survival are distinct from those that influence dendritic spine formation.

## Results

### Dsn1 and Ndc80 deficiency trigger progressive neuronal death

Dsn1 is an essential component of the Mis12 complex of the kinetochore and we had previously generated mice carrying a floxed allele of Dsn1 with which to study the role of the protein in dendritic spine formation [5]. While characterizing the dendritic phenotypes associated with Dsn1 deficiency, we noted that hippocampal cultures did not survive well after deletion of Dsn1. Primary hippocampal neurons from *Dsn1^fl/fl^* mice were infected with lentivirus expressing either active Cre or an inactive Cre (ΔCre) that served as a control. These constructs were expressed under the neuron-specific synapsin promoter and fused with a nuclear localization signal (NLS) and GFP to ensure localized expression and fluorometric identification of infected neurons. Eleven days post-infection, cell viability was assessed using propidium iodide (PI) staining and determining the fraction of GFP-expressing cells that were positive for PI. We observed a significant increase in neuronal mortality in Dsn1-deficient neurons (∼70%) compared to control cultures (∼10%) (Figure 1A, B, and D). This lethal phenotype was effectively rescued by the viral re-expression of Dsn1, which reduced cell death to approximately 15% (Figures 1C, C’ and D). The requirement of Dsn1 for neuronal survival was not limited to hippocampal neurons; the same increase in PI positive cells was observed upon Dsn1 deletion from cultures of cortical neurons (Fig S1A-D).

**Figure 1.**
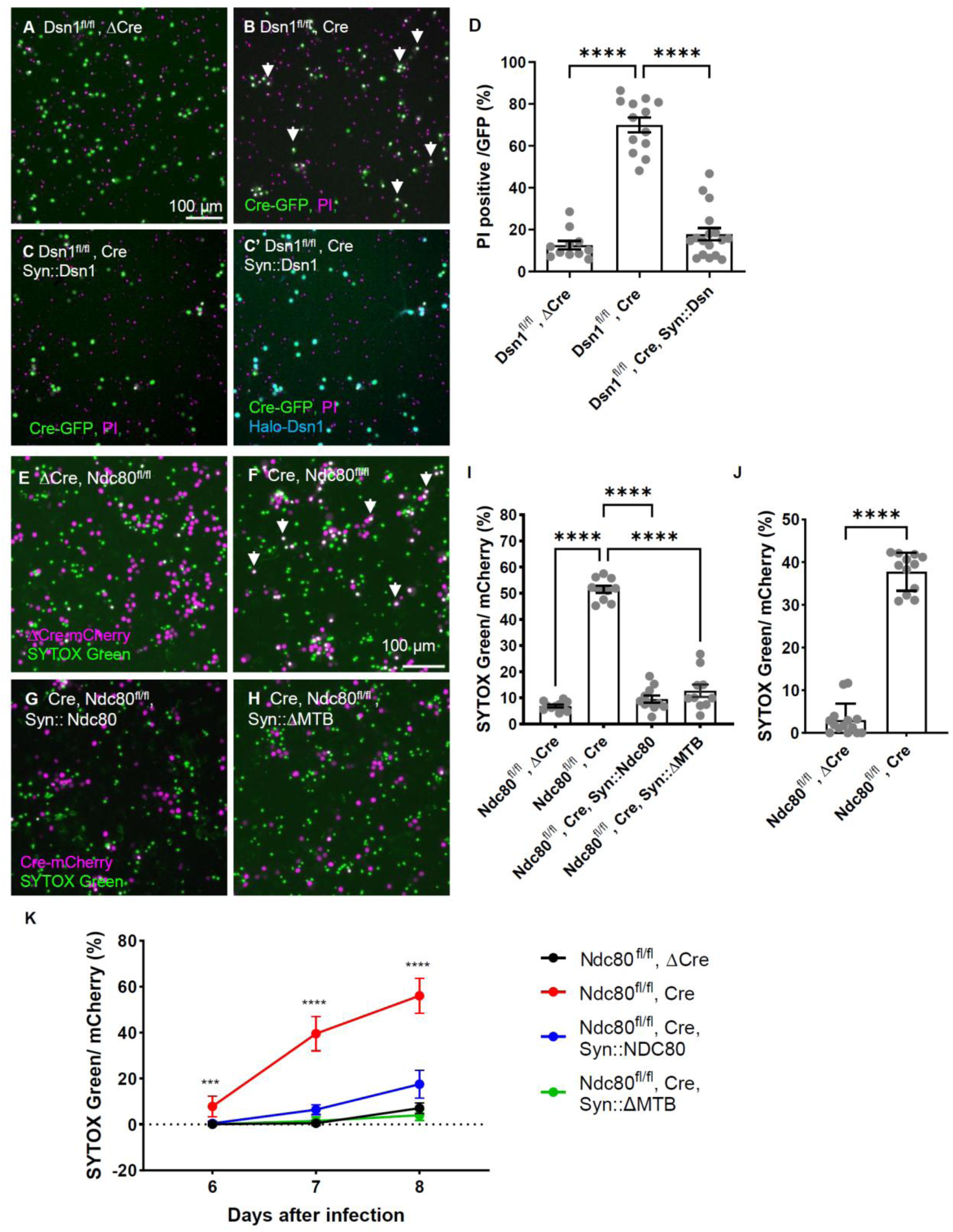
Loss of Dsn1 or Ndc80 *in vitro* induces neuronal death. **(A–C)** Representative fluorescence images of *Dsn1^fl/fl^* hippocampal neurons at day *in vitro* (DIV) 14. Neurons were infected on DIV 3 with lentiviruses expressing: **(A)** ΔCre-GFP (control), **(B)** Cre-GFP (knockout), or **(C, C’)** Cre-GFP with Halo-Dsn1 (rescue). Cell death was visualized using the membrane-impermeant nucleic acid stain propidium iodide (PI). **(D)** Quantification of the percentage of GFP-expressing neurons that are PI-positive (non-viable) from images as in (A-C). **(E-H)** Representative fluorescence images of *Ndc80^fl/fl^* hippocampal neurons on DIV 9. Neurons were infected on DIV 1 with lentiviruses expressing: **(E)** ΔCre-mCherry (control), **(F)** Cre-mCherry (knockout), **(G)** Cre-mCherry with Halo-NDC80 (wild-type rescue), or **(H)** Cre-mCherry with Halo-ΔMTB-NDC80 (microtubule-binding deficient rescue). Cell death was visualized using the membrane-impermeant nucleic acid stain SYTOX Green. **(I)** Quantitative analysis of images as in (E-H) of the percentage of mCherry-expressing neurons that are SYTOX Green-positive (non-viable) across the indicated experimental conditions. **(J)** Quantification, similar to (I) of cell death of *Ndc80^fl/fl^* cortical neurons infected on DIV 6 with virus expressing either active CRE-mCherry or inactive ΔCre-mCherry and imaged on DIV13. **(K)** Quantification of dying hippocampal neurons over time following Cre-mediated Ndc80 excision, from images as in Fig S2A-D. The graph depicts the percentage of mCherry-expressing cells that are SYTOX Green-positive at 6, 7, and 8 days post-expression of Cre or ΔCre and in cultures where neuronal viability was rescued by re-expression of Halo-Ndc80 or Halo-ΔMTB-NDC80, Error bars represent mean ± SEM. Statistical significance was determined by one-way ANOVA with Tukey’s post-hoc test; ****p < 0.0001.

We examined the time course of neuronal decline in hippocampal cultures by a similar protocol, but with SYTOX Green, a high-affinity nucleic acid stain that selectively penetrates compromised plasma membranes, as the marker of dying cells. Already at 7 days post-infection with the CRE-expressing virus, the percentage of mCherry-positive cells that were also SYTOX Green positive was significantly increased. The percentage of dying cells increase further at 9 days and at each time point could be rescued by expression of Dsn1 (Fig S1E-H).

To determine if other kinetochore proteins were also required for neuronal survival, we used hippocampal cultures from our previously generated *Ndc80^fl/fl^* mice [5]. Ndc80 is the kinetochore component that directly interacts with microtubules in mitosis [1, 3]. We performed the same SYTOX Green assay described above, again infecting cells via lentivirus expressing a Cre-recombinase construct or an inactive Cre (ΔCre) and assaying their viability 8 days later. Approximately 50% of CRE-infected neurons were SYTOX-positive, compared to only 7% in control cultures (Figure 1 E, F, and I). As observed for Dsn1, loss of Ndc80 in cultures of cortical neurons also increased counts of dying cells (Fig 1J).

### The microtubule-binding function of Ndc80 is not required for preserving neuronal viability

Because Ndc80 is the microtubule binding subunit of the kinetochore [3, 10, 13],and because the ability to bind microtubules is required for its regulation of dendritic microtubules and spine formation [5], we hypothesized that failure to bind microtubules would also account for the cell death phenotype in neuronal cultures. To our surprise, this was not the case. The microtubule binding region of Ndc80 has been mapped and a deletion of 80 amino acids from the N-terminus of Ndc80 prevents this interaction [14, 15]. We previously confirmed that this deletion (ΔMTB), with 3 additional mutations of glutamate residues also important for microtubule binding, prevented the colocalization of expressed Ndc80 with microtubule plus ends in the neuronal cytoplasm and prevented the ability of Ndc80 to restore plus-end dynamics [5]. We therefore attempted to rescue the viability of hippocampal neurons after deletion of Ndc80 by viral expression of either wild-type Ndc80 or the microtubule-binding deficient variant (ΔMTB). We quantified the percentage of dying infected cells with SYTOX green 6, 7, and 8 days after infection. In marked contrast to what we had observed for the dendritic phenotypes, both constructs could restore neuronal viability at each time point (Figures 1 G, H, I and K and S2 A-E). Thus, the microtubule-binding property of Ndc80 is dispensable for neuronal survival, and the function of Ndc80 in preserving neuronal viability is distinct from its action in regulating dendrite development.

### Nuclear Dsn1 is likely required to preserve neuronal viability

Our previous findings indicated that kinetochore proteins in neurons localize to both the nucleus and the cytosol of cell bodies and neurites [5]. To identify the subcellular compartment where Dsn1 exerts its pro-survival function, we generated N-terminal Halo-tagged Dsn1 constructs fused with either a nuclear localization signal (NLS) or a nuclear export signal (NES). These variants were expressed under the synapsin promoter to evaluate their respective rescue efficiencies in Dsn1-knockout neurons (Figure 2 A-G). To test the compartmentalization of these constructs, we performed live-cell imaging using Halo ligands (Figure 2A, B), While NLS-Halo-Dsn1 was sequestered within the nucleus and undetectable in the cytoplasm, NES-Halo-Dsn1 localized more broadly. Although present in the cytosol of the soma and neurites, NES-Halo-Dsn1 also accumulated in the nucleus, likely due to suboptimal NES-mediated export in these cells.

**Figure 2.**
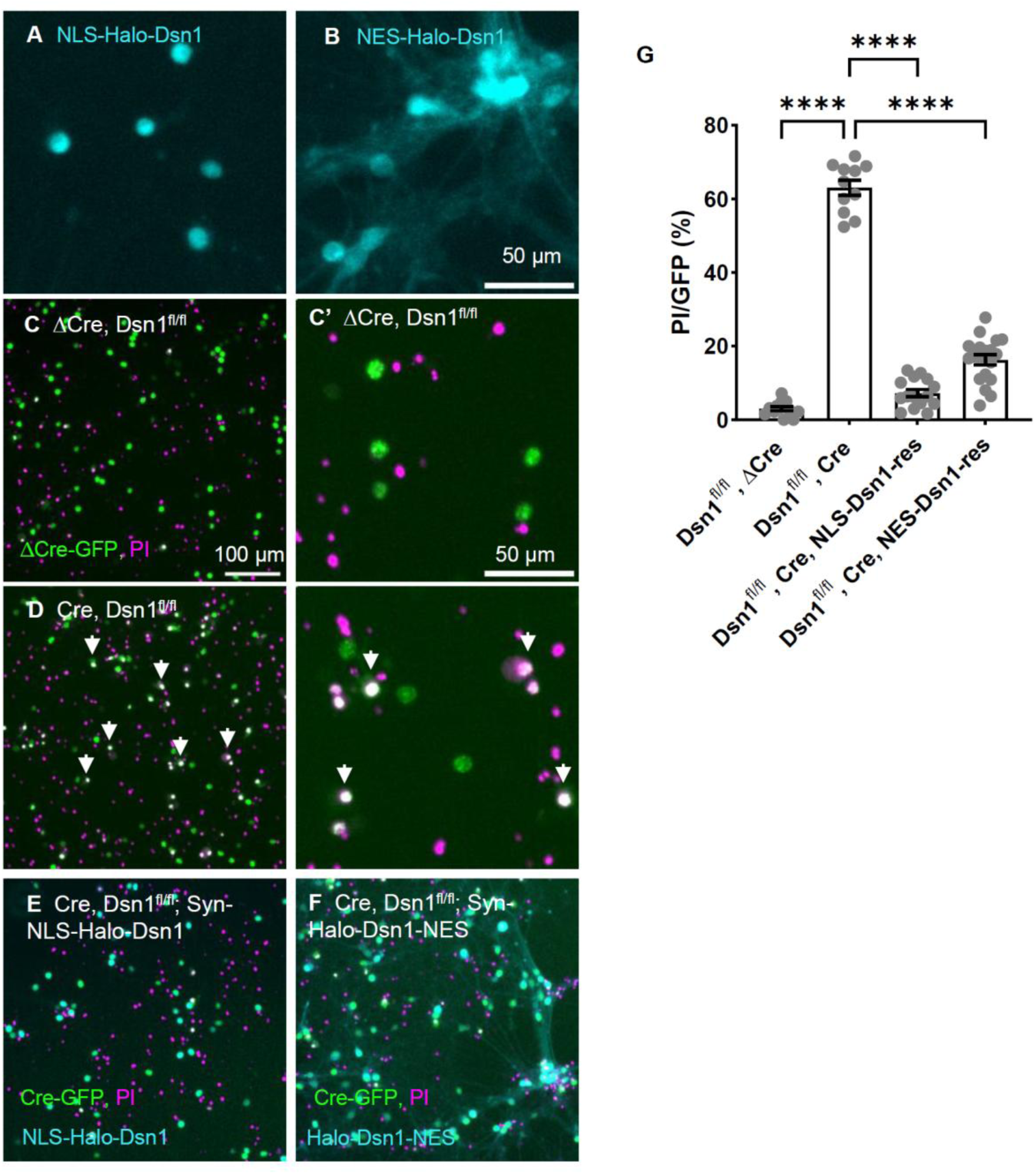
Nuclear Dsn1 is essential for neuronal survival. **(A-B)** Localization of NLS-Halo-Dsn1 **(A)** and NES-Halo-Dsn1 **(B)** in hippocampal neurons. Whereas the NLS-tagged construct appears to be confined to the nucleus, the NES-tagged construct is present in both nucleus and cytoplasm. **(C–F)** Evaluation of the ability of NLS– and NES-tagged constructs to rescue the viability of DIV 13 neurons deleted for Dsn1. *Dsn1^fl/fl^* neurons were infected on DIV 1 with either **(C)** ΔCre-GFP, or **(D)** Cre-GFP alone or together with **(E)** NLS-Halo-Dsn1 or **(F)** NES-Halo-Dsn1. **(C’ and D’)** Enlarged detail of panels C and D showing ΔCre-GFP (C’) or Cre-GFP (D’) expressing neurons stained with cell death marker PI. **(G)** Quantification of images as in (C-F) of the percent of GFP-positive neurons also positive for PI, demonstrating rescue efficiency of both Dsn1 variants. Data represents the mean ± SEM; ****p < 0.0001 by one-way ANOVA with Tukey’s post-hoc test.

Both NLS-Halo-Dsn1 and NES-Halo-Dsn1 significantly mitigated the cell death phenotype caused by deletion of Dsn1 (Figure 2E, F and G). NLS-Dsn1 proved somewhat superior in its capacity to rescue survival compared to the NES variant. That the highly nuclear-localized variant was most effective at rescue, and that the rescuing NES variant also was present in nuclei, strongly suggests that a nuclear function of Dsn1 is required to maintain neuronal viability and that the loss of this nuclear function is responsible for the death of neurons upon Dsn1 deletion.

### Ndc80 or Dsn1 deficiency triggers neuronal apoptosis via DNA damage

To characterize the mode of cell death induced by Ndc80 or Dsn1 removal, we assessed markers of apoptosis after CRE expression in hippocampal cultures of the floxed alleles. Immunostaining revealed a significant increase in cleaved caspase-3-positive neurons in Dsn1 and Ndc80 cultures after CRE expression compared to controls (Figure 3A-I). Consistent with our viability data, this apoptotic phenotype was fully rescued by the exogenous expression of either Dsn1 (for the *Dsn1^fl/fl^* cultures) or either wild-type Ndc80 or the ΔMTB microtubule-binding deficient variant (for the *Ndc80^fl/fl^* cultures) (Figure 3A-I). These findings emphasize that while Ndc80’s microtubule-binding capacity is essential for dendritic spine maintenance [5], it is dispensable for neuronal survival. To further test the significance of this apoptotic pathway for the death of the hippocampal neurons, we undertook pharmacological inhibition of caspases using Quinoline-Val-Asp-Difluorophenoxymethylketone (Q-VD-Oph). Caspase inhibition significantly attenuated Ndc80-deficient cell death (Figure 3J and S3), confirming that these neurons undergo canonical apoptosis.

**Figure 3.**
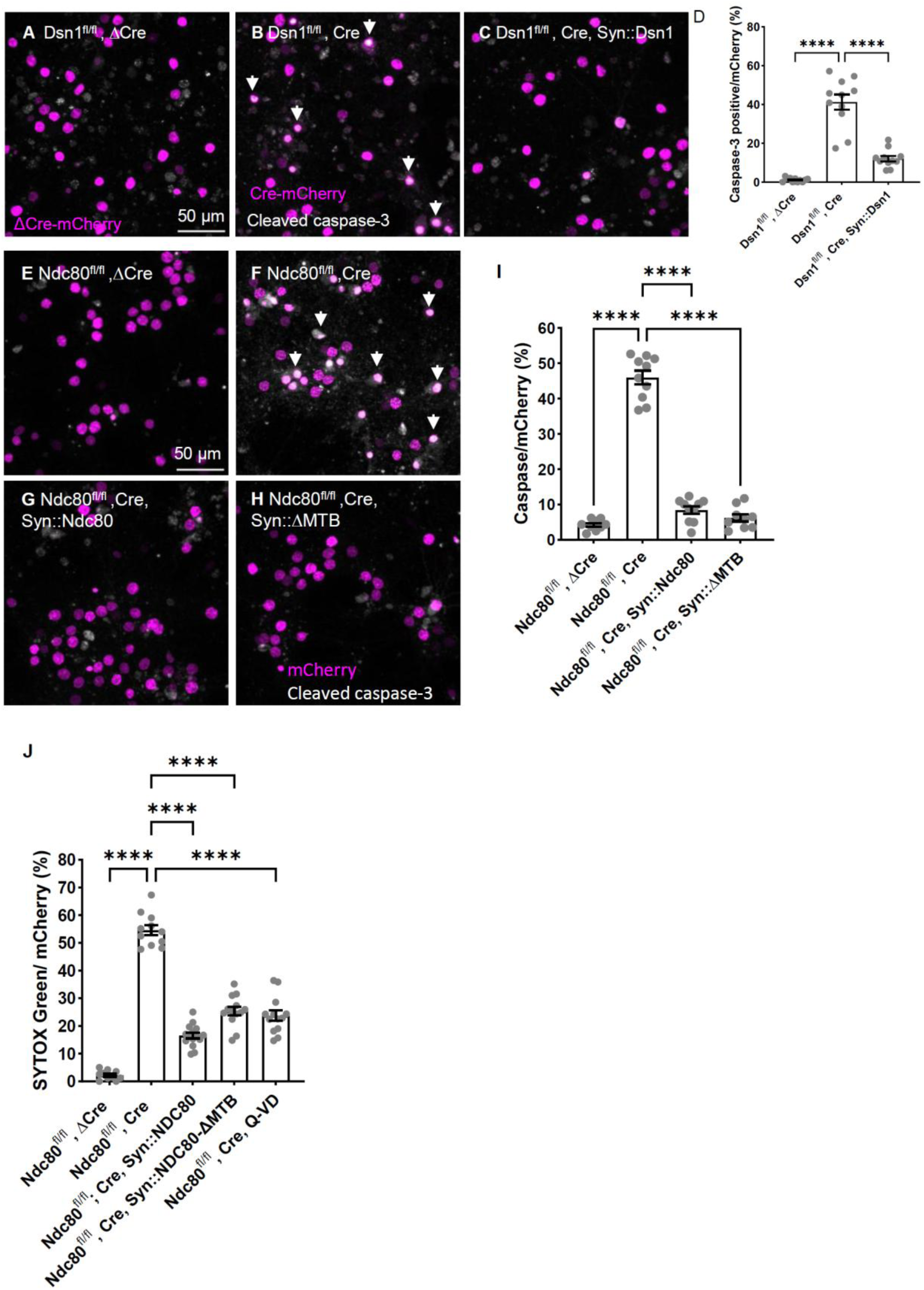
Ndc80 and Dsn1 deficiency triggers neuronal apoptosis. **(A–C)** Representative images of DIV 19 *Dsn1^fl/fl^* hippocampal neurons infected on DIV 9 with the indicated lentiviral constructs. Cultures were immunostained for the apoptotic marker cleaved caspase-3. **(D)** Quantification of the percentage of mCherry-positive neurons that are cleaved caspase-3 positive from images as in A-C. Data represents the mean ± SEM. Statistical significance was determined by one-way ANOVA with Tukey’s post-hoc test; ***p < 0.0001. **(E–H)** Representative images of DIV 10 *Ndc80^fl/fl^* hippocampal neurons infected on DIV 1 with indicated mCherry-tagged lentiviral constructs. Cultures were immunostained for the apoptotic marker cleaved caspase-3. **(I)** Quantification of the percentage of mCherry-positive neurons that are cleaved caspase-3 positive from images as in E-H. Data represents the mean ± SEM. Statistical significance was determined by one-way ANOVA with Tukey’s post-hoc test; ****p < 0.0001. **(J)** Quantification of the percentage of mCherry positive neurons that are CYTOX green positive from images as in figure S3 A-E. Statistical significance was determined by one-way ANOVA with Tukey’s post-hoc test; ****p < 0.0001.

Biochemical analysis corroborated these results, showing elevated levels of both cleaved caspase-3 and cleaved PARP in lysates of cultures from which Dsn1 or Ndc80 had been deleted (Figure 4A, B). Given that DNA double-strand breaks are potent triggers of apoptosis and an increase in PARP cleavage can be a consequence of extensive DNA damage[16], we examined γ-H2AX foci, a hallmark of double-strand breaks. Quantification revealed that after Ndc80 removal, 87% of neurons exhibited five or more γ-H2AX foci, compared to 7% in controls (Figure 4C-E). Similarly, Dsn1 removal caused a significant accumulation of γ-H2AX foci (84% vs 6%), which was largely reversed upon re-expression of Dsn1 under the synapsin promoter (Figure 4F-I).

**Figure 4.**
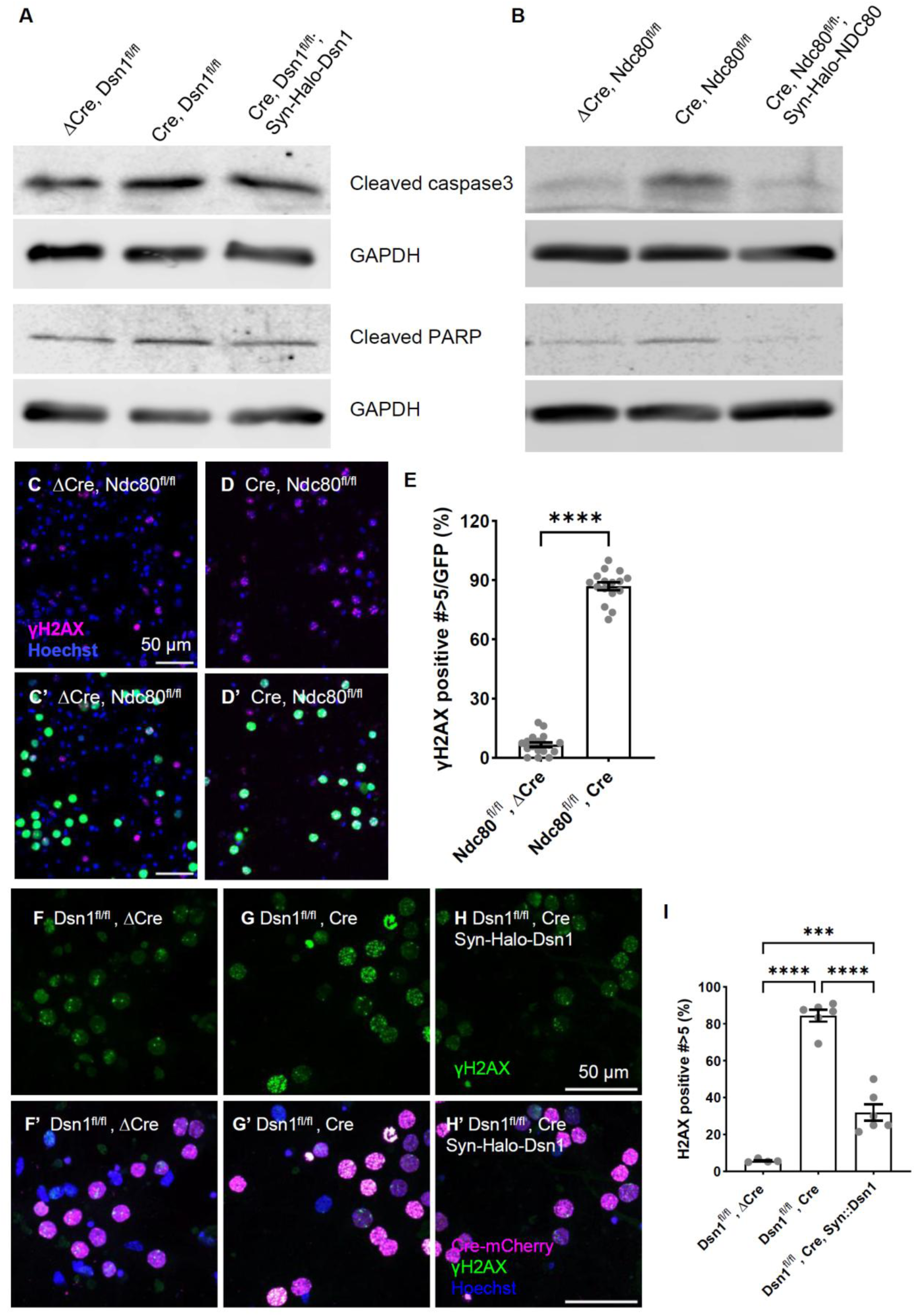
Dsn1 and Ndc80 loss induce DNA double strand breaks. **(A and B)** Western blot indicating increased cleaved caspase-3 and cleaved PARP levels in *Dsn1^fl/fl^* (**A**) and *Ndc80^fl/fl^* (**B**) neurons after Cre expression and its prevention by re-expression of Dsn1 or Ndc80. **(C and D)** Immunofluorescence images of γ-H2AX staining (magenta) in *Ndc80^fl/fl^* after expression of Cre-GFP or ΔCre-GFP. Nuclei are marked with Hoechst 33342 (blue) and in **(C’ and D’)** the GFP channel (green) is added to mark the infected cells. **(E)** Quantification from images as in C’ and D’ of the percentage of GFP-positive neurons with ≥5 ã-H2AX Data represents the mean ± SEM. foci/nucleus. Statistical significance was determined by Unpaired Welch’s t Test ****p < 0.0001. **(F-H, F’-H’)** Immunofluorescence images of γ-H2AX staining (green) in in *Dsn1^fl/fl^* neurons after expression of control ΔCre-mCherry (**F**), Cre-mCherry (**G**) and Cre-GFP with re-expression of Dsn1 (**H**). In (**F’-H’**) the mCherry and nuclear markers are added to indicate nuclei expressing the Cre and ΔCre constructs. **(I)** Quantification of the percentage of Cre-mCherry-positive neurons with ≥5 γ-H2AX foci/nucleus, highlighting the rescue of DNA damage upon Dsn1 re-expression. Data represent the mean ± SEM. Statistical significance was determined by one-way ANOVA with Tukey’s post-hoc test; ****p < 0.0001.

### Kinetochore components are essential for RGC survival *in vivo*

To determine whether Ndc80 and Dsn1 were also important for neuronal survival *in vivo*, we targeted Retinal Ganglion Cells (RGCs) of *Ndc80^fl/fl^* mice by intravitreal injections of AAV-Cre-GFP or AAV-ΔCre-GFP under control of the synapsin promoter. Age-matched (littermate) pairs of mice were injected with the Cre and ΔCre constructs and tissue was collected from pairs at 38, 49, 69, 78, 88, 94, 108, 111 or 132 days post-injection. In retinal flat-mounts, loss of Ndc80 reduced RGC density (RBPMS-positive cells) relative to the ΔCre control from that pair (Figures 5 and Supplemental data). This was apparent in every pair examined. For a more detailed statistical analysis, we employed a linear mixed effects model to combine data from these timepoints and to consider the possibility that the interval between intravitreal injection and tissue collection influenced the extent of cell loss (see Supplemental Data). In aggregate the effect size was approximately a 40% reduction in RGC density after 111 days of infection and this was independent of the interval before tissue collection over the range examined. Similarly, we assessed the impact of Dsn1 loss by injecting AAV-Cre-GFP into *Dsn1^fl/fl^* mice and collecting tissue, 40, 90, 110, and 114 days post-injection, Dsn1-KO retinas also exhibited significant RGC loss, retaining just 35% of control RGCs 114 days post injection (Figure 6 and Supplemental Data). Collectively, these data indicate that the kinetochore KMN complex operates as a functional unit essential for the long-term survival of neurons *in vivo* and that loss of kinetochore components even from adult mice will result in gradual neuronal loss.

**Figure 5.**
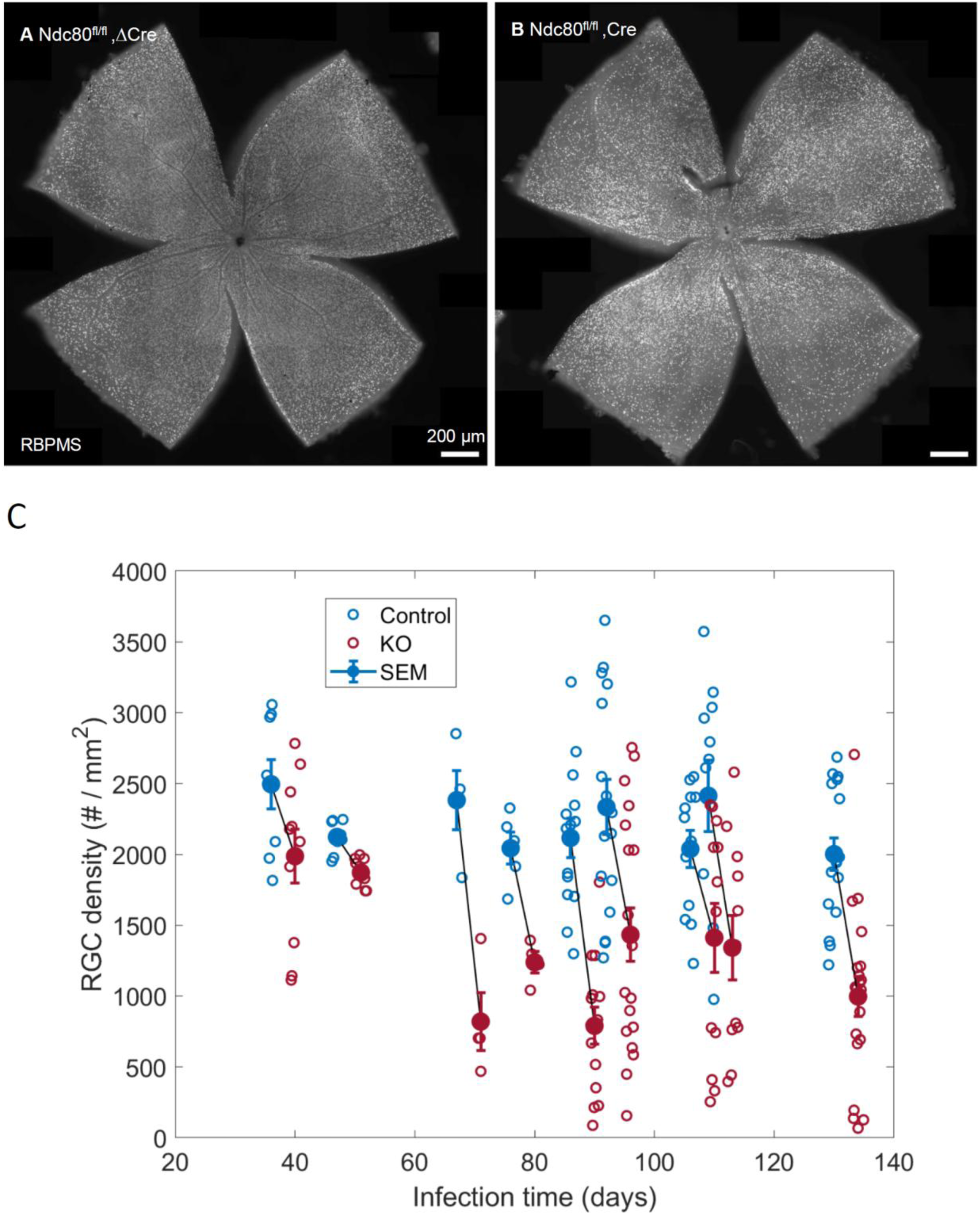
In vivo Ndc80 depletion results in significant RGC attrition. **(A–B)** Flat-mount retinal images of adult *Ndc80^fl/fl^* mice injected with AAV-ΔCre-GFP (A, control) or AAV-Cre-GFP (B, KO) at postnatal day 51 (P51) and analyzed at P162. Retinas were stained for the RGC-specific marker RBPMS. **(C)** Effect of Ndc80 deletion on RGC density. For each set of littermate mice connected by a black line, one mouse received an intravitreal injection of CRE expressing virus (red data points) and one received the inactive CRE (blue). Open circles reflect separate retinal areas from a mouse; closed circles indicate the mean value of RGC density for that mouse. The pairs are ordered along the x-axis according to duration between the time of viral injection and tissue collection (infection time). All pairs show a clear loss of RGCs upon CRE expression regardless of the infection time over this range, although it is less pronounced at the shortest times. For more details on the statistical analysis, see Supplemental Methods.

**Figure 6.**
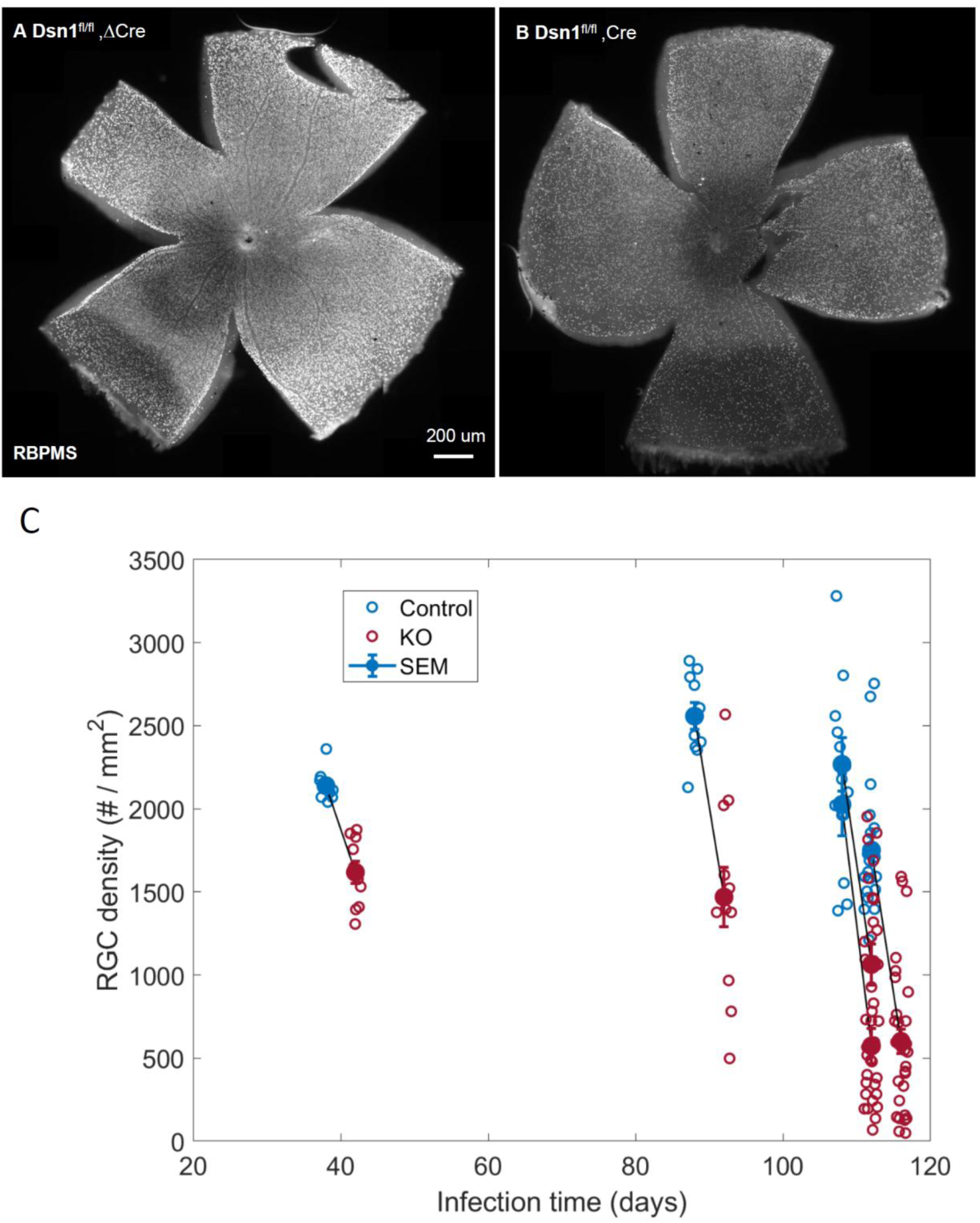
Dsn1 is required for long-term RGC survival in vivo. **(A–B)** Representative images of flat-mount retinal images of *Dsn1^fl/fl^* mice injected with AAV-ΔCre-GFP (**A**, control) or AAV-Cre-GFP (**B**) at postnatal day 1 and analyzed at P115. Retinas are stained for RGC-specific marker RBPMS. **(C)** Effect of Dsn1 deletion on RGC density. For each set of littermate mice connected by a black line, one mouse received an intravitreal injection of CRE expressing virus (red circles) and one received the inactive CRE (blue circles). Open circles reflect separate retinal areas from a given mouse; closed circles indicate the mean value of RGC density for that mouse. The pairs are ordered along the x-axis according to duration between the time of viral injection and tissue collection (infection time). All pairs show a clear loss of RGCs upon CRE expression regardless of the infection time over this range. For more details on the statistical analysis, see Supplemental Methods.

## Discussion

In this study, we have identified an essential role for proteins of the outer kinetochore complex—specifically Ndc80 and Dsn1—in maintaining the survival of post-mitotic neurons. While these proteins are canonical components of the mitotic apparatus for chromosome segregation, our findings demonstrate that their depletion in differentiated neurons triggers a progressive apoptotic program. This neurodegeneration is characterized by the accumulation of DNA double-strand breaks (DSBs) and the activation of caspase-dependent death pathways. Inhibition of caspases prevented neuronal death. The phenotypes reported here, as well as previously reported dendritic phenotypes[4, 5], are not restricted to developing neurons or neurons regenerating in culture. Delivery of CRE-expressing virus to the vitreous space and consequent deletion of Ndc80 or Dsn1 resulted in a profound loss of retinal ganglion cells regardless of the developmental stage at which the knockout was induced. This highlights that these kinetochore protein’s non-mitotic role in survival continues even into adulthood. We note, however, that cell death proceeded substantially slower in vivo than in culture.

Previous genetic studies of kinetochore proteins in postmitotic neurons pointed strongly to the presence of a complex very much like the mitotic kinetochore. In *Drosophila* and *C. elegans*, we and others found that components of each of the subcomplexes of the mitotic kinetochore gave equivalent neuronal phenotypes, whether scored at neuromuscular junctions, in sensory dendrites, or in positioning of sensory neurons [4–6, 8]. In addition, we found that loss of one component changed the distribution of another, suggesting they functioned as a complex[4]. In mammalian neurons, we have only examined three elements of the outer kinetochore, but these include both core components (Mis12 and Dsn1) and the crucial microtubule-binding subunit (Ndc80). In the present study as well as our previous study of dendritic microtubules and spines[5], these yield similar phenotypes when deleted. Moreover, we showed by PLA reaction that Mis12 and Ndc80 are in close proximity to one another in both neurites and cell bodies[5]. Thus, the neuronal pro-survival function reported here likely also involves action of a kinetochore-like complex. A biochemical characterization of the neuronal kinetochore, however, is lacking and aspects of its composition may differ from the mitotic complex.

### The neuronal kinetochore functions differently in promoting survival and in limiting dendritic spine formation

Though Dsn1 and Ndc80 are now implicated in two aspects of post-mitotic neurons, the mechanisms by which they promote survival and restrict dendritic spine formation are distinct. The difference is most clearly established by ΔMTB, the Ndc80 construct that lacks its microtubule-binding domain. This construct rescued neuronal survival (Figures 1 and S2) although microtubule-binding was essential for rescue of the dendritic phenotype. In this regard, the requirement of Ndc80 for neuronal survival is extraordinary; Ndc80 has no other known function that does not entail its interactions with microtubules. In mitosis, these are the microtubules of the spindle apparatus and in postmitotic neurons they are the microtubules of axons and dendrites. In dendrites, as in mitosis, the kinetochore is associated with microtubule plus ends and limits the numbers of actively growing plus ends[5]. Herein lies another distinction between the dendritic and pro-survival functions of these proteins. Whereas the kinetochore was functioning in the cytoplasm to associate with and limit the dynamics of plus ends, our data pointed to a strictly nuclear role for Dsn1 in suppressing apoptosis. The Dsn1 protein with a nuclear localization signal (NLS-Halo-Dsn1) appeared strictly localized to nuclei and yet rescued neuronal viability in the Dsn1 deleted neurons. Indeed, it rescued better than the NES-Halo-Dsn1 construct that was present in both nuclei and cytosol (Figure 2). Though we cannot exclude the presence of NLS-Halo-Dsn1 in the cytosol below the level of detection, the weight of evidence favors a nuclear requirement.

The kinetochore is commonly viewed as a load-bearing linker that binds to microtubules at one end and chromatin at the other. Concerning the cytoplasmic kinetochore function in neurons, one end clearly interacts with microtubules, but, as there is no chromatin in the dendrites, it is unknown what, if anything interacts with the other end. In contrast, with regard to the apparently nuclear kinetochore function reported here, it is possible that the proteins are connected to chromatin, but whether there are interactions at the other end is unknown. Indeed, it is hard to speculate as to the cell-survival function of Ndc80, because it’s microtubule-binding function was unnecessary. For neither of the neuronal functions of these proteins is a role for load-bearing and tension-sensing apparent, two kinetochore properties that are crucial in mitosis.

### Post-mitotic kinetochore and cell survival

How does the loss of Dsn1 or Ndc80 activate an apoptotic cascade in which DNA damage causes PARP activation and caspase cleavage? Potentially, the connection is quite indirect, involving changes to transcription that only indirectly feedback to trigger apoptosis. More direct connections are, however, conceivable. One possibility is that the complex is required for proper chromosome organization, and, absent the kinetochore, chromosomes are more susceptible to breakage. Alternatively, the kinetochore proteins may be part of the DNA damage response and, in their absence, strand breakage goes unrepaired with the consequence of persistent γH2AX and eventual cell death. This scenario might explain why neurons die more slowly in vivo than in culture, where the stress of dissociation may induce DNA damage. Support for this model is suggested by the known role of components of the kinetochore-associated Spindle Assembly Complex (Bub1 and Mad1) in the DNA Damage Response[17, 18].

An intriguing question that remains outstanding concerns the regulation of kinetochore proteins in neurons. In dividing cells, Ndc80, Dsn1, and the close partner of Dsn1, Mis12, are each the target of phosphorylations by the kinases that regulate progression through the cell cycle. Accurate regulation of the kinetochore is essential for guaranteeing chromosome alignment at the midline and proper segregation to the daughter cells. Future studies may elucidate parallel avenues of regulation for the neuronal kinetochore and its role in the survival of postmitotic neurons.

## Supporting information

Method details

Key resource table

## Acknowledgements

We thank Lala Mkhitaryan for technical support. This work was supported by NIH grant R37NS109211 to TLS. We would like to acknowledge the Cellular Imaging and Gene Manipulation Cores of the IDDRC at Boston Children’s Hospital (NIH P50 HD105351) and the Viral Core (NIH P35 EY012196). GZ was supported by NIH Developmental Neurology training grant T32NS007473.

## Author Contributions

G.Z. and T.L.S. conceived of the project and G.Z. performed most of the experiments. H.M, Q.W. and F.T., under the guidance of Z.H., conducted some *in vivo* viral injections. C.D. supplied iRGCs. R.T.B. provided statistical analysis, G.Z. and T.L.S. wrote the manuscript, with contributions from all co-authors.

## Supplemental figure and legends

**Figure S1.**
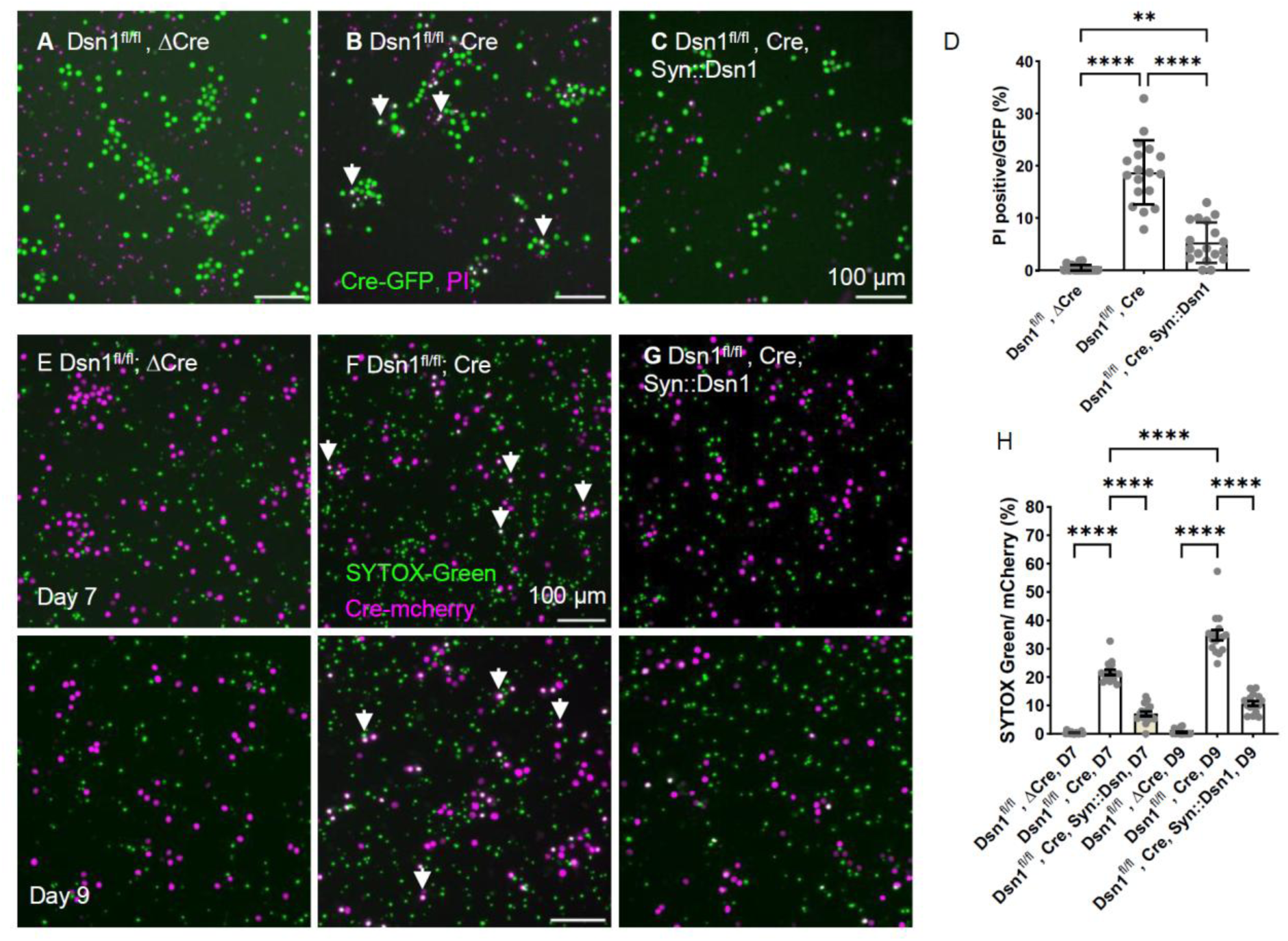
Loss of Dsn1 causes progressive neuronal death. **(A-C)** Representative images of *Dsn1^fl/fl^* cortical neurons infected with lentiviruses expressing: (**A**) ΔCre-GFP (control), (**B**) Cre-GFP (knockout), or (**C**) Cre-GFP with Halo-Dsn1 (rescue). Cell death was visualized using the membrane-impermeant nucleic acid stain propidium iodide (PI, magenta). Neurons were infected on DIV 9 and imaged on DIV 16. **(D)** Quantification of the percentage of GFP-expressing neurons (green) that are PI-positive (non-viable, white) from images as in (**A-C**). Statistical significance was determined by one-way ANOVA with Tukey’s post-hoc test; ****p < 0.0001. **(E-G)** Representative images showing the temporal increase in dying mCherry-expressing cells (white) in Dsn1 knockout (**F**), rescue (**G**) cultures compared to controls (**E**). (**H**) Quantification of dying cells from images like those in E-G. Note the significant escalation in neuronal attrition beginning at day 7 and greater at day 9 post infection. Viability could be rescued at each time point by restoration of Dsn1 expression. Statistical significance was determined by one-way ANOVA with Tukey’s post-hoc test; ****p < 0.0001.

**Figure S2.**
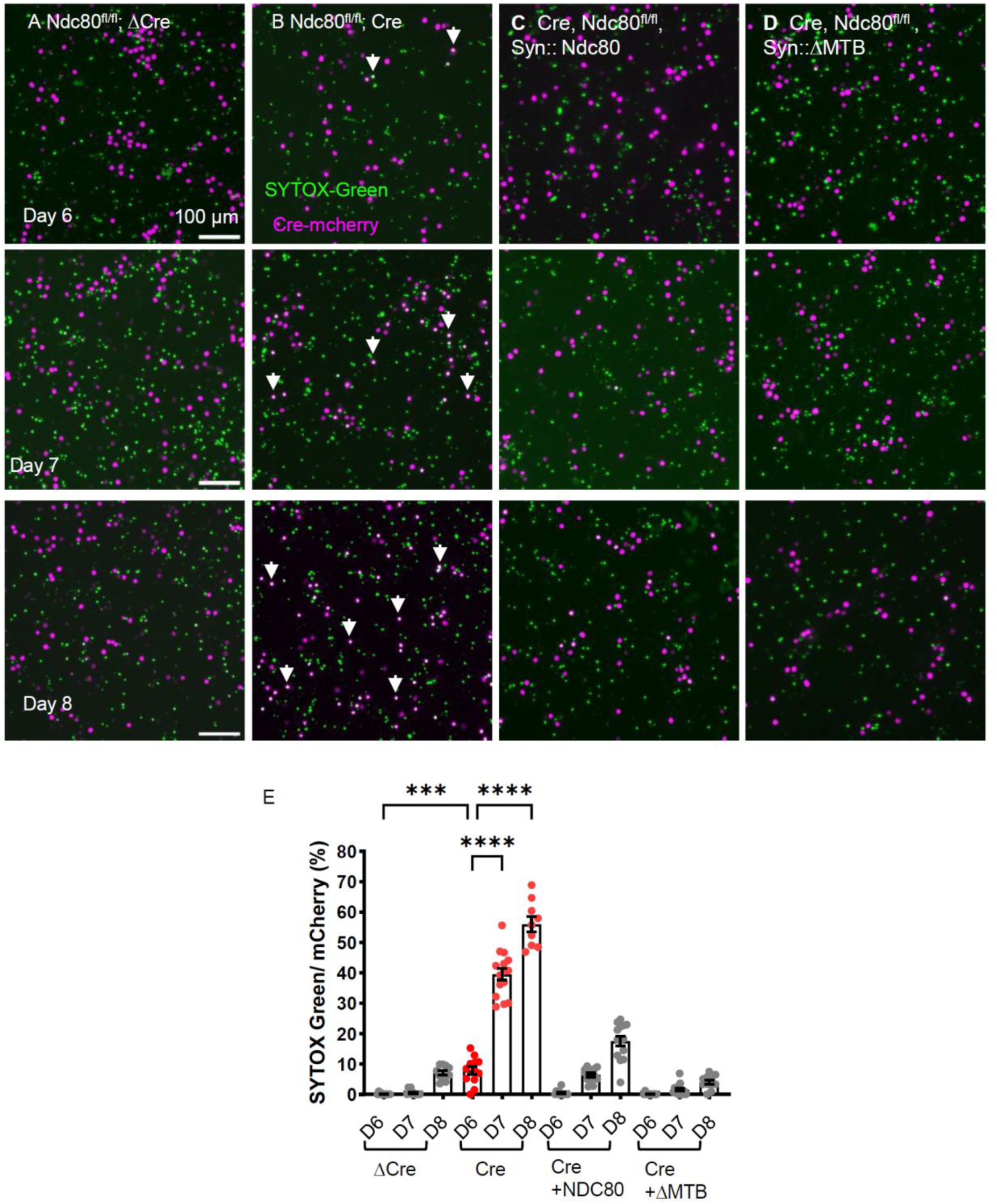
Loss of Ndc80 causes progressive neuronal death. (**A-D**) Progression of neuronal death in Ndc80^fl/fl^ hippocampal neurons at 6, 7, and 8 days post-expression of mCherry-tagged ΔCre (**A**) Cre (**B**), Cre and NDC80 (**C**), or Cre and ΔMTB (**D**). (**E**) Quantification of the percentage of SYTOX Green-positive Cre-expressing neurons from images as in A-D. The data are replotted as a function of time in Figure 1K. Data represents the mean ± SEM. Statistical significance was determined by one-way ANOVA with Tukey’s post-hoc test; ***p < 0.0001.

**Figure S3.**
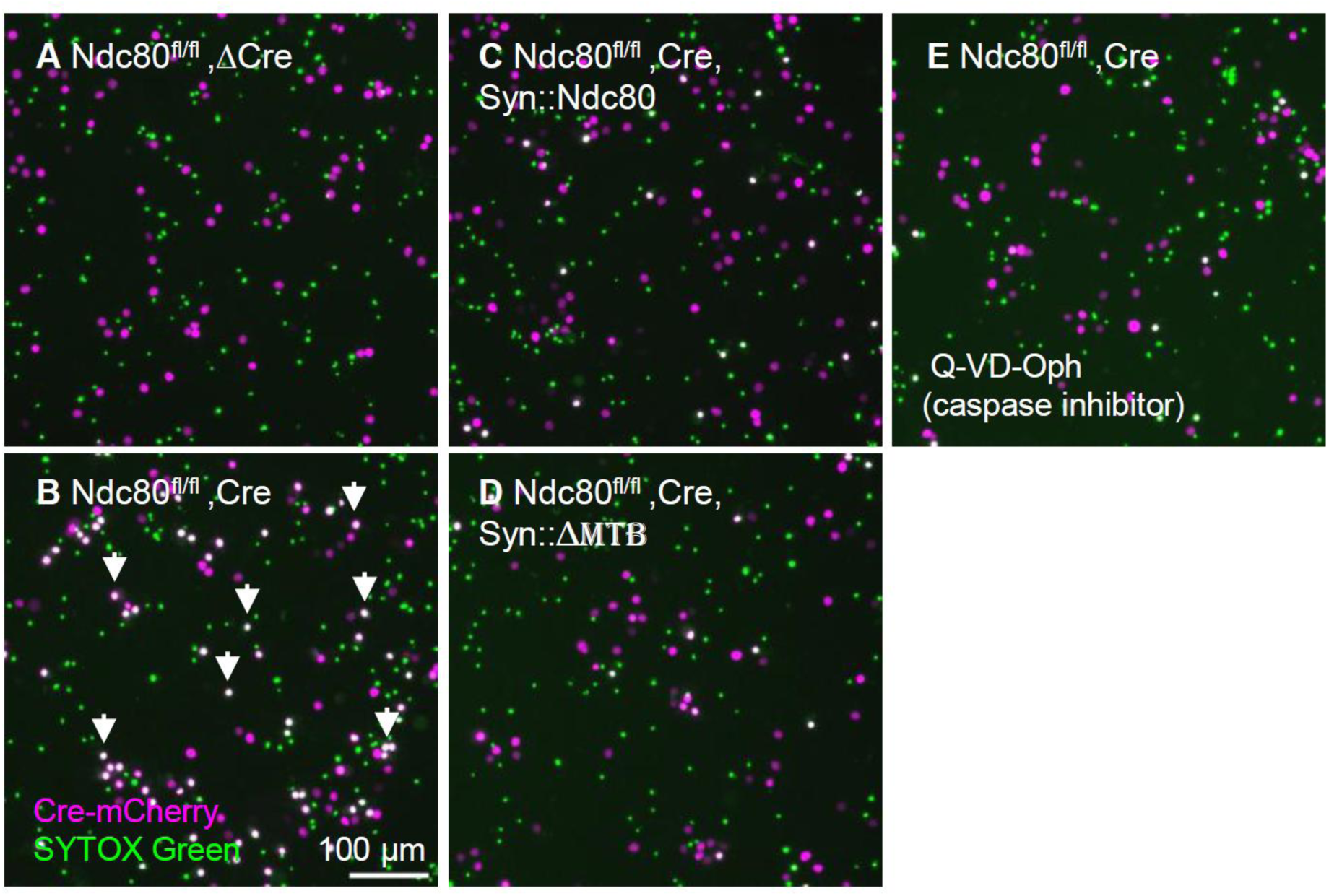
Caspase inhibition attenuates Ndc80-deficient neuronal death. **(A–E)** Representative images of *Ndc80^fl/fl^* hippocampal neurons at days in vitro (DIV) 17, following lentiviral infection on DIV 9 with: (**A**) ΔCre-mCherry (control), (**B**) Cre-mCherry (knockout), (**C**) Cre-mCherry and Halo-NDC80 (rescue), or (**D**) Cre-mCherry and Halo-ΔMTB-NDC80 (microtubule-binding deficient rescue), or (**E**) infected with Cre-mCherry on DIV 9 and treated with the pan-caspase inhibitor Q-VD-Oph on DIV 11 to assess apoptotic blockade. All neurons were stained with the membrane-impermeant cell death marker SYTOX Green.

**Figure S4.**
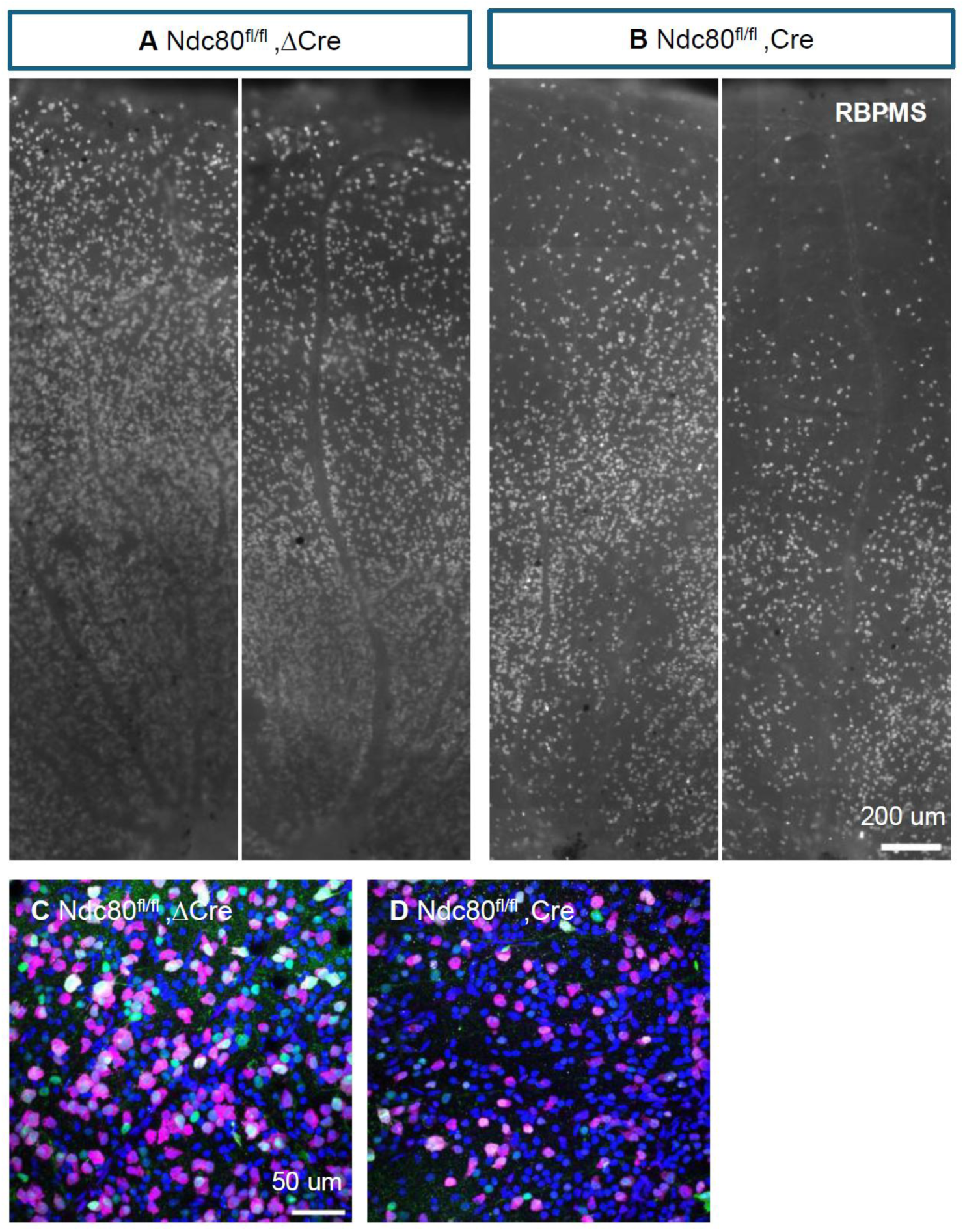
Timing of Ndc80 deletion does not alter RGC loss severity. **(A–B)** Representative images of retinal flat-mounts from *Ndc80^fl/fl^* mice injected with AAV-ΔCre-GFP (control) or AAV-Cre-GFP (knockout) at postnatal day 32 (P32) and sacrificed at P126. RGCs were identified via immunostaining for the specific marker RBPMS. **(C–D)** Magnified views of the regions from (A) and (B). The GFP signal confirms successful viral transduction and expression of ΔCre or active Cre.

