## Supplementary material for "The kinetochore proteins Ndc80 and Dsn1 are required for survival of postmitotic neurons": Method details

#### Plasmids

pLXV-hSyn-Halo-DSN1, pLXV-hSyn-Halo-NDC80, pLXV-hSyn-Halo-ΔMTB-NDC80, AAV2-hSyn-Cre-2A-EGFP and AAV2-hSyn-ΔCre-2A-EGFP were constructed using the Gibson assembly (NEB, E2611) method as decribed in [1].

#### AAV Virus production

For Cre expression, we used a plasmid derived from AAV:ITR-U6-sgRNA(backbone)-hSyn-Cre-2A-EGFP-KASH-WPRE-shortPA-ITR (Addgene, 60231) from which the KASH had been removed. A recombination-deficient version of Cre (ΔCre) was used as control [2]. These plasmids were used by Boston Children's Hospital Viral Core to prepare 7m8 serotype AAV for Cre or ΔCre expression. Then, these viruses were used for transducing retinal ganglion cells in vivo. The titers of all viral preparations were at least $5 \times 10^{12}$ genome copies/ml.

#### Lentivirus production

pFSW-hSyn-NLS-Cre-mCherry, pFSW-hSyn-NLS-ΔCre-mCherry, pFSW-hSyn-NLS-Cre-GFP, pFSW-hSyn-NLS-ΔCre-GFP, pLXV-hSyn-Halo-DSN1, pLXV-hSyn-Halo-NDC80, and pLXV-hSyn-Halo-ΔMTB-NDC80 plasmids were packaged in lentivirus by co-transfecting HEK293 cells with plasmid psPAX2 (Addgene, 12260) and a VSV-G envelope expressing plasmid (either pMD2.G; Addgene). The viral particles were purified by centrifugation and resuspended in DPBS+0.001% F68 (Thermofisher, Cat#24040032). All viral particles were flash-freezing in liquid nitrogen and storage at -80°C.

#### Generation of conditional knockout mice

The Dsn1 and Ndc80 conditional knockout mice were generated by the Gene Manipulation Core at Boston Children's Hospital as described in [1].

#### Intravitreous AAV injection

Intravitreal virus injection was performed at age between P1 to P50. Briefly, a pulled-glass micropipette was inserted near peripheral retina behind the ora serrata and deliberately angled to avoid damage to the lens. 2 ul of AAV2/7m8-hSyn-Cre-GFP or AAV2/7m8-hSyn-ΔCre-GFP virus were injected for mice [3]. The titter of each virus was adjusted to 7 x $10^{12}$ genome copies/mL.

**Western blot for detecting the cleaved caspase3 and cleaved PARP**

Primary mouse cortical neurons were seeded at a density of 1x$10^6$ cells per well of 6 well plates and transduced at DIV2 with a lentiviral vector expressing either Cre-mCherry or ΔCre-mCherry under the control of a synapsin promoter. The rescue wells co-transduced with Cre-mCherry and Halo-DSN1 or Halo-NDC80 under the synapsin promoter. Seven days post-infection, neurons were harvested and lysed in 120 μL/well of RIPA buffer supplemented with a 1:100 dilution of protease inhibitors. Lysates were incubated on ice for 30 minutes, followed by centrifugation at 12,000 rpm for 10 minutes at 4°C. The resulting supernatants were collected, combined with 40 μL of 4× Laemmli sample buffer, and denatured by boiling for 10 minutes. Following equilibration to room temperature, 30 μL of each sample was resolved via 12% SDS-PAGE and analyzed by Western blotting. Target proteins were detected using rabbit anti-cleaved caspase3 (Cell signaling, 9664, 1:1000), rabbit anti-Cleaved PARP1 (Cell signaling 954, 1:1000) and mouse anti-GAPDH (Millipore Sigma, CB1001, 1:2000) primary antibodies.

**Immunohistochemistry of mouse retina**

Mice were euthanized with > 5% isoflurane for 5 min and then perfused with cold PBS and 4% PFA/PBS. Dissected eyes were immersed in 4% PFA/PBS for post-fixation 1 hour at room temperature. Eyes were washed with PBS 4 times, transferred to PBS and dissected the retina out under microscope. The following staining was performed in 96 well plate. Washed twice in PBS and permeabilized with 1% Triton-X100 (PBST) for at least 2 hours. retinas were then washed twice with PBS before blocking with SuperBlock buffer (ThermoFisher, Cat # 37515) for 1 hour. All washes were 10-15 minutes. retinas were incubated with primary antibodies in blocking buffer for 2 overnights at 4°C. After 5 washes with PBS, they were incubated with secondary antibodies in blocking buffer for overnight at 4°C. After 5 washes with PBS, retina was mounted in Fluoromount-G Anti-Fade mounting medium (SouthernBiotech, cat# 0100-35).

We used chicken anti-GFP (1:1000; Aveslab, GFP-1010), guinea pig anti-RBPMS (Novus, NBP2-80389) and rabbit anti-phosphorylated γ-H2AX (Cell signaling, 9718) as primary antibodies, and Alexa Fluor-488, 568 and 647 conjugated antibody (1:400; ThermoFisher Scientific) as secondary antibody. The retinas were imaged with an LSM700 laser-scanning confocal microscope (Zeiss) and a 40× 1.4 NA objective using separate channels, at least 6 fields (100 um from the periphery of retina) from each retina were imaged and processed using the Fiji (ImageJ).

**Analysis of cell death in cultured neurons**

Hippocampal neurons were cultured as in [4]. Specifically, hippocampal neurons were obtained from E18 mixed-sex mouse embryonic brains, plated on 24 well glass bottom plates (Cellvis, cat# P24-1.5H-N) coated with 20 µg/mL poly-L-Lysine (Sigma-Aldrich, P2636) and 3.5 µg/mL laminin (ThermoFisher Scientific, 23017015). The neurons were maintained in Neurobasal medium (ThermoFisher Scientific, 21103049) supplemented with B27 (ThermoFisher Scientific, 17504044), L-glutamine (ThermoFisher Scientific, 25030081), and penicillin/streptomycin (Sigma, p4333).

Cultured hippocampal neurons were infected between DIV1 to DIV10 with lentivirus pFSW-hSyn-ΔCre-mCherry (control) or pFSW-hSyn-Cre-mCherry, or pFSW-hSyn-ΔCre-GFP (control) or pFSW-hSyn-Cre-GFP and, when indicated, also with pLXV-hSyn-Halo-Ndc80, pLXV-hSyn-Halo-ΔMTB-Ndc80, or pLXV-hSyn-Halo-Dsn1 for rescue. After 6 to 12 days infection, neurons were incubated with 150 nM CYTOX Green (Fishersci, S7020) or 500nM Propidium iodide (PI) (ThermoFisher Scientific, P1304MP) for 15 minutes, then take the images under Nikon TiEclipse Inverted Microscope with 10x object. For immunostaining, the neurons were fixed for 12 min with 4% paraformaldehyde, 4% sucrose in phosphate-buffered saline at room temperature. After 3x 10-minute washes with PBS and blocking with Superblock buffer for 30 minutes, cells were immunostained in Superblock blocking buffer (ThermoFisher, 37515) overnight at 4℃ with rabbit anti- Cleaved caspase3 antibody (Cell signaling, 9664) or rabbit anti-γ-H2AX (Cell signaling, 9718) at 1: 800, and subsequently incubated with goat anti-rabbit Alexa-488 or goat anti-rabbit Alexa-568 secondary antibody (1:500 in Superblock) for 3 hours at room temperature. The stained samples were mounted with cover slips using Fluoromount-G® Anti-Fade mounting medium (SouthernBiotech, 0100-35). Z-stack images were acquired with the same setting for same batch of samples under Zeiss LSM700 confocal microscope (Zeiss, Thornwood, NY) for quantification the density of cleaved caspase3 and cleaved PARP. All neurons that express GFP or mCherry (which marked the Cre or inactive Cre expressing cells) were imaged in an unbiased manner. To quantify the signal density, neurons were selected based on GFP or mCherry signal as not to bias the selection. For each phenotype, at least 4 fields were captured. the images were acquired with a 40X, 1.4 NA oil immersion objective with 0.5 µm intervals and

1024x1024 pixel resolution and averaged twice. The cleaved caspase3 and cleaved PARP signal were then analyzed manually by using Fiji ImageJ software (available at https://fiji.sc/).

### QUANTIFICATION AND STATISTICAL ANALYSIS

#### Statistical analysis

Statistical analysis of data was done using GraphPad Prism software. P values were derived using a student's t-test with Welch-correction or One-way ANOVA with Tukey's multiple comparisons test as indicated. Graphs were plotted in GraphPad Prism using the scatter dot plot functions showing the mean and standard error of the mean (SEM). P values for each experiment are indicated in the figures or figure legends.

For dead neuron quantification, the neurons which mCherry positive (represents express Cre or ∆Cre) and both mCherry positive and CYTOX Green positive were counted respectively from each field, the percentage of dead neurons were calculated by (both mCherry and CYTOX Green positive neuron # / mCherry positive neuron #) * 100. Similarly, the percentage of dead neurons which were infected with GFP-Cre were calculated by (both GFP and PI positive neuron # / GFP positive neuron #) * 100. The percentage of caspase3 positive neurons were calculated by (both cleaved caspase3 and mCherry positive neuron # / mCherry positive neuron #) * 100. For γ-H2AX staining quantification, since the γ-H2AX has basic staining signal for almost all neurons, we made an arbitrary cut for 5 foci of γ-H2AX as the standard. The percentage of γ-H2AX positive neurons were calculated by (both GFP and the foci number of γ-H2AX above 5 neuron # / GFP positive neuron #)*100. At least four fields were counted for each condition.

For RGC death quantification, the RBPMS positive cell number were counted from every field of each retina, the RGC density were calculated by RBPMS positive cell #/ image area ($mm^2$). Here are the details about the analysis.

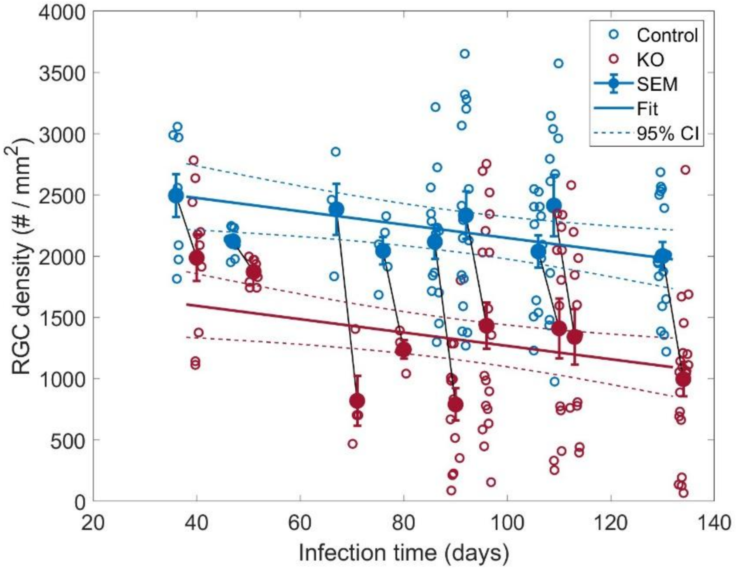

*Figure 1.* Ndc80 KO experiment. Open circles represent the raw data of RGC counts from each of multiple fields-of-view normalized by retinal area for all animals. The filled circles indicate the mean ± SEM for each animal; thin black lines connect animals that were paired, each from the same litter and having the same survival time post-infection. Thick colored lines show the fits of the linear mixed effect model (fixed effects only; see Supplemental Methods) along with the 95% confidence intervals (dashed lines).

The data shown in Figure 1 are hierarchical and "nested": within each litter, multiple animals were tested and "paired" with respect to survival time; and multiple technical replicates (open circles) were taken from each animal's retinas. This nested design creates dependencies in the data which require a corresponding multi-level analysis[5]. This was done using a mixed-effects linear regression model[6,7], with fixed effects for the main covariates, namely Cre treatment and infection time, and random effects to account for the nested design and repeated measures. In Wilkinson notation[8] the full model was:

RGCdensity ~ 1 + rx + infectionTime + (1 | litterID) + (1 | litterID:mouseID) +
(1 | litterID:mouseID:fieldNum) (1)

where *RGCdensity* is the RGC count normalized by the area of the field-of-view, *rx* is an indicator variable set to '1' for mice that received Cre and '0' for control animals, *infectionTime* represents the survival time after infection, and the terms in parentheses are the random effects as follows:

(1 | litterID) accounts for the variance between different litters,
(1 | litterID:mouseID) uniquely identifies a mouse within its specific litter, and
(1 | litterID:mouseID:fieldNum) treats 'fieldNum' (individual RGC counts) as a random intercept grouping factor to account for the repeated measurements from single mice.

Although 'infection time' was not systematically manipulated in these experiments, including it as a fixed effect in the model allowed us to better estimate the treatment effect while accounting for the fact that different pairs of animals had different survival times. The full model was fit to the data with the 'fitlme' function in MATLAB (MathWorks, Natick, MA), using maximum likelihood. The results are shown in **Table 1**.

**Table 1**. Results for full model (1). Model AIC = 3128.5; adjusted $R^2$ = 0.98

| Name | Estimate | Std. Error | t-stat | DF | p-Value | 95% CI (lower) | 95% CI (upper) |
|---|---|---|---|---|---|---|---|
| y-intercept | 2693.31 | 202.82 | 13.28 | 195 | $4.38 \times 10^{-29}$ | 2293.31 | 3093.31 |
| rx | -880.34 | 119.44 | -7.37 | 195 | $4.71 \times 10^{-12}$ | -1115.89 | -644.79 |
| infectionTime | -5.45 | 2.05 | -2.65 | 195 | 0.0086 | -9.49 | -1.40 |

All fixed parameters were statistically significant ($p < 0.05$), and the model accounted for 98% of the variance, after adjusting for the number of parameters. The magnitude of the *rx* parameter tells us that, on average, across all animal-pairs and infection time-points, the retinas treated with Cre had 880 ± 119 fewer RGCs per $mm^2$. The negative value of the *infectionTime* parameter indicates that there was a progressive loss of RGCs over time in both groups, though this effect was small (Cohen's d-prime = 0.19).

To give a more intuitive interpretation of the effect size, for the average mouse with an 80-day survival time (middle of the range in **Figure 1**), the model predicts an RGC density of 2,258 in controls and 1,377 in treated animals, amounting to a 39% reduction in RGC density due the KO of  NDC80.

Results were similar for the animals in which Dsn1 was knocked out (**Figure 2**), although here the effect of infection time was slightly more pronounced (Cohen's d-prime = 0.32).

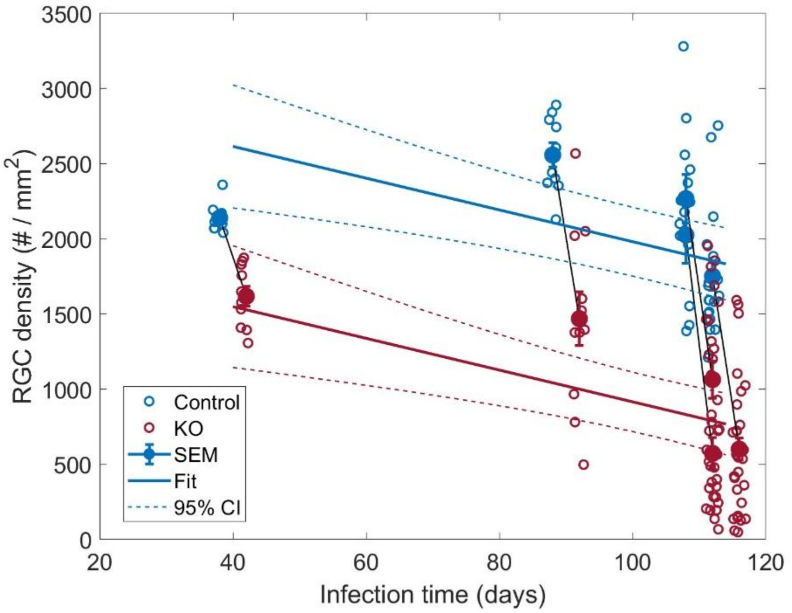


*Figure 2.* Dsn1 KO experiment. Open circles represent the raw data of RGC counts from each of multiple fields-of-view normalized by retinal area for all animals. The filled circles indicate the mean ± SEM for each animal; thin black lines connect animals that were paired, each from the same litter and having the same survival time post-infection. Thick colored lines show the fits of

the linear mixed effect model (fixed effects only; see Supplemental Methods) along with the 95% confidence intervals (dashed lines).

The same regression model fit to these data yielded the results shown in **Table 2**.

**Table 2**. Results for model (1) fit to Dsn1 data. Model AIC = 2186.7; adjusted $R^2$ = 0.98

| Name | Est. | SE | tStat | DF | pValue | Lower | Upper |
|---|---|---|---|---|---|---|---|
| y-intercept | 3037.43 | 306.28 | 9.92 | 141 | $6.51 \times 10^{-18}$ | 2431.93 | 3642.92 |
| rx | -1065.48 | 129.84 | -8.21 | 141 | $1.29 \times 10^{-13}$ | -1322.16 | -808.79 |
| infectionTime | -10.56 | 2.80 | -3.77 | 141 | $2.41 \times 10^{-04}$ | -16.10 | -5.02 |

For the average mouse with a 70-day survival time (middle of the range in **Figure 2**), the model predicts an RGC density of 2,298 in controls and 1,233 in treated animals, amounting to a 46% reduction in RGC density due to KO of Dsn1.

For both data sets, we ran an additional model including an interaction term for *rx* and *infectionTime*, which would allow for the possibility that there was a more (or less) rapid loss of RGCs in the Cre-treated animals over time. However, in neither case was this term statistically significant (Ndc80, p = 0.161; Dsn1, p = 0.057), so we omitted this term from the final model that is shown in Figures 1 and 2.

Finally, we performed two diagnostic tests of the models fit to each data set, which, taken together, indicate that our use of the linear model was appropriate. A plot of the raw residuals vs. the fitted values (**Figure 3 & 4**, left) shows reasonable homoscedasticity and linearity, while the Q-Q plot displays the approximate normality of the residuals (**Figures 3 & 4**, right).

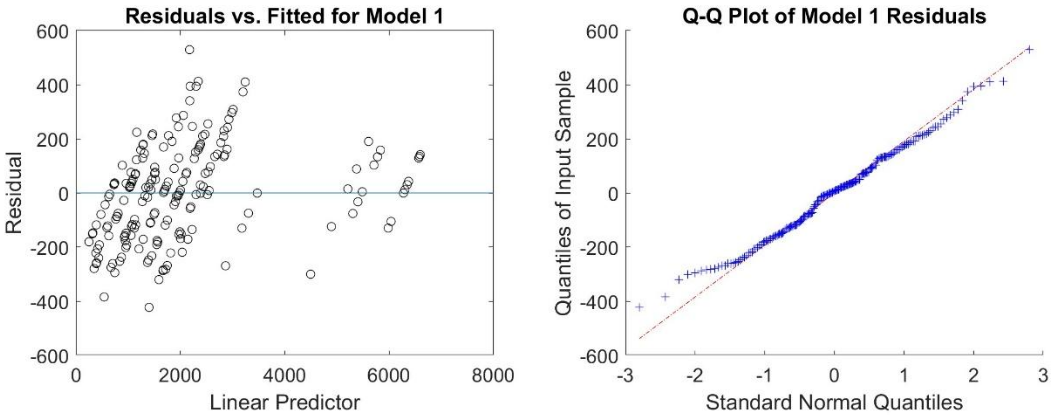


*Figure 3.* Regression diagnostics for model (1) fit to the Ndc80 data.

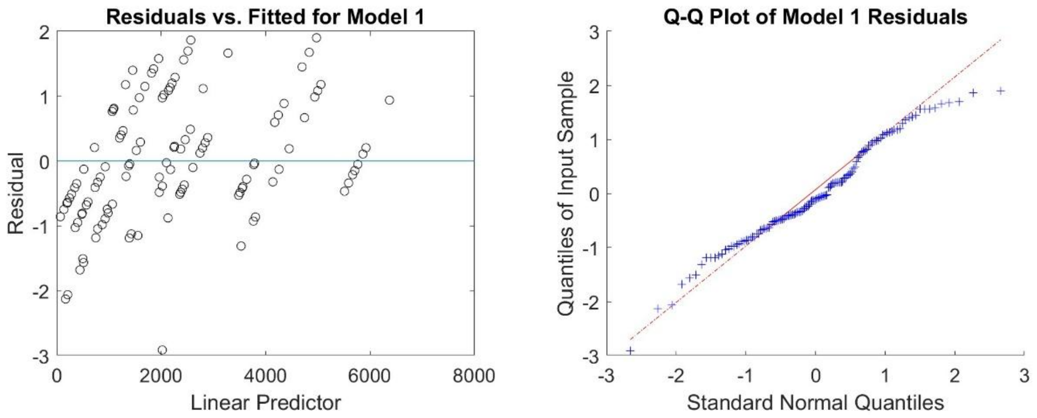


*Figure 4.* Regression diagnostics for model (1) fit to the Dsn1 data.

The raw data, analysis code and results are available at:
https://github.com/rickborn/RGC_Survival-paper


1. Zhao, G., Sharma, A., Tang, J., Aleman, M., Liang, X., Miner, L., Qi, J., Xiang, W., Tian, F., Goldberg, Y., et al. (2026). Kinetochore proteins control microtubule dynamics in postmitotic neurons to regulate the formation of dendritic spines. Proc Natl Acad Sci U S A *123*, e2520684123. 10.1073/pnas.2520684123.
2. Pollina, E.A., Gilliam, D.T., Landau, A.T., Lin, C., Pajarillo, N., Davis, C.P., Harmin, D.A., Yap, E.L., Vogel, I.R., Griffith, E.C., et al. (2023). A NPAS4-NuA4 complex couples synaptic activity to DNA repair. Nature *614*, 732–741. 10.1038/s41586-023-05711-7.
3. Tian, F., Cheng, Y., Zhou, S., Wang, Q., Monavarfeshani, A., Gao, K., Jiang, W., Kawaguchi, R., Wang, Q., Tang, M., et al. (2024). Core transcription programs controlling injury-induced neurodegeneration of retinal ganglion cells. Neuron *112*, 2453–2456. 10.1016/j.neuron.2024.06.009.
4. Zhao, G., Oztan, A., Ye, Y., and Schwarz, T.L. (2019). Kinetochore Proteins Have a Post-Mitotic Function in Neurodevelopment. Dev Cell. 10.1016/j.devcel.2019.02.003.
5. Aarts, E., Verhage, M., Veenvliet, J.V., Dolan, C.V., and van der Sluis, S. (2014). A solution to dependency: using multilevel analysis to accommodate nested data. Nat Neurosci *17*, 491–496. 10.1038/nn.3648.
6. Laird, N.M., and Ware, J.H. (1982). Random-effects models for longitudinal data. Biometrics *38*, 963–974.
7. Fitzmaurice, G.M., Laird, N.M., and Ware, J.H. (2011). Applied longitudinal analysis, 2nd Edition (Wiley).
8. Wilkinson, G.N., C. E. Rogers (1973). Symbolic Description of Factorial Models for Analysis of Variance. Applied Statistics *22*, 392–399. https://doi.org/10.2307/2346786.
