## Supplementary material for "The kinetochore proteins Ndc80 and Dsn1 are required for survival of postmitotic neurons": Key resource table

### KEY RESOURCES TABLE

| REAGENT or RESOURCE | SOURCE | IDENTIFIER |
|---|---|---|
| **Antibodies** | | |
| rabbit anti-cleaved caspase3 | Cell signaling | 9664<br>RRID: **AB_2070042** |
| Rabbit anti-Cleaved PARP | Cell signaling | 9544<br>RRID:AB_2160724 |
| Chicken anti-GFP | Aveslabs | GFP-1010<br>RRID:AB_10000240 |
| Rabbit anti-γ-H2AX | Cell signaling | 9718<br>RRID:AB_2118009 |
| Guinea pig anti-RBPMS | Novus | NBP2-80389<br>RRID:AB_3409455 |
| Mouse anti-GAPDH (6C5) | Millipore Sigma | CB1001<br>RRID:AB_2107426 |
| Goat anti-Chicken IgY (H+L) Secondary Antibody, Alexa Fluor™ 488 | Thermofisher | A-11039 |
| Phalloidin–Atto 647N | Millipore Sigma | 65906 |
| Goat anti-Rabbit IgG (H+L) Highly Cross-Adsorbed Secondary Antibody, Alexa Fluor™ 568 | Thermofisher | A-11036 |
| Alexa Fluor® 647 AffiniPure® F(ab')$_2$ Fragment Donkey Anti-Guinea Pig IgG (H+L) | Jackson immuonResearch | 706-606-148 |
| Goat anti-Rabbit IgG (H+L) Highly Cross-Adsorbed Secondary Antibody, Alexa Fluor™ 488 | Thermofisher | A-11034 |
| **Critical Commercial Assays** | | |
| **Experimental Models: Cell Lines** | | |
| HEK293T | ATCC | CRL-3216<br>RRID: CVCL_0063 |
| **Experimental Models: Organisms/Strains** | | |
| *Dsn1 floxed mouse* | Schwarz lab [1] | |
| *Ndc80 floxed mouse* | | |
| **Oligonucleotides** | | |
| **Recombinant DNA** | | |
| pLXV-hSyn-Halo-Ndc80 | [1] | N/A |
| pLXV-hSyn-Halo-ΔMTB-Ndc80 | [1] | N/A |
| pLXV-hSyn-Halo-Ndc80-3KE | [1] | N/A |
| pLXV-hSyn-Halo-Dsn1 | [1] | N/A |

| | | |
|---|---|---|
| AAV: hSyn-Cre-2A-EGFP | [1] | |
| AAV: hSyn-ΔCre-2A-EGFP | [1] | |
| pFSW-hSyn-NLS-Cre-GFP | Michael Greenberg lab | |
| pFSW-hSyn-NLS-ΔCre-GFP | Michael Greenberg lab | |
| pFSW-hSyn-NLS-Cre-mCherry | Michael Greenberg lab | |
| pFSW-hSyn-NLS-ΔCre-mCherry | Michael Greenberg lab | |
| **Software and Algorithms** | | |
| Fiji | Schindelin, J.; Arganda-Carreras, I. & Frise, E. et al. (2012) | https://fiji.sc/ |
| Adobe illustrator | adobe | https://www.adobe.com/products/illustrator.html |
| GraphPad Prism | GraphPad Software | https://www.graphpad.com/scientific-software/prism/ |
| Other | | https://www.zeiss.com/microscopy/int/confocal.html |
| LSM700 laser-scanning confocal microscope | Zeiss | https://www.micro-shop.zeiss.com/?p=us&f=o&a=v&m=s&id=440762-9904-000 |
| 63× 1.4 NA objective | Zeiss | https://www.leica-microsystems.com/products/confocal-microscopes/p/leica-tcs-sp8/ |
| Nikon TiEclipse Inverted Microscope | Nikon | |

1. Zhao, G., Sharma, A., Tang, J., Aleman, M., Liang, X., Miner, L., Qi, J., Xiang, W., Tian, F., Goldberg, Y., et al. (2026). Kinetochore proteins control microtubule dynamics in postmitotic neurons to regulate the formation of dendritic spines. Proc Natl Acad Sci U S A *123*, e2520684123. 10.1073/pnas.2520684123.
